# Hedgehog produced by the Drosophila wing imaginal disc induces distinct expression responses in three target tissues

**DOI:** 10.1101/2020.03.05.979799

**Authors:** Ryo Hatori, Thomas B. Kornberg

## Abstract

Hedgehog (Hh) is an evolutionarily conserved signaling protein that has essential roles in animal development and homeostasis. We investigated Hh signaling in the region of the Drosophila wing imaginal disc that produces Hh and is near the tracheal air sac primordium (ASP) and myoblasts. Hh distributes in concentration gradients in the wing disc anterior compartment, ASP, and myoblasts and activates different sets of genes in each tissue. Some transcriptional targets of Hh signal transduction are common to the disc, ASP, and myoblasts, whereas others are tissue-specific. Signaling in the three tissues is cytoneme-mediated and cytoneme-dependent. We conclude that a single source of Hh in the wing disc regulates cell type-specific responses in three discreet target tissues.

**Summary:** Hedgehog produced by the wing imaginal disc signals to wing disc, myoblast and tracheal cells

## Introduction

Morphogens are evolutionarily conserved signaling proteins that pattern tissues during development and regulate cell differentiation in stem cell niches. They spread across the tissues they target, generating distributions that elicit concentration-dependent responses. The Hh morphogen was discovered because of its roles in embryonic and imaginal disc development (Mohler, 1988; Nüsslein-Volhard and Wieschaus, 1980), and in humans, defects in Hh signaling have been associated with congenital diseases and have been implicated in malignancies, including basal cell carcinoma, medulloblastoma, and pancreatic cancer (Hui and Angers, 2011; Roberts et al., 2017). Elucidating the function and mechanisms of Hh signaling is important to both developmental biology and medicine.

Studies of Hh signaling in the wing pouch primordium of the Drosophila wing imaginal disc have elucidated many fundamental features of Hh signaling. The wing disc is a flattened sac with two closely juxtaposed and connected single-cell layered sheets. The sheet with columnar epithelial cells contains most of the disc cells and includes the progenitors of both the wing and notum. In the wing primordium, *hh* is expressed exclusively in the posterior compartment cells and Hh protein disperses to form a concentration gradient that extends over a distance of approximately ten anterior compartment cells. In a concentration dependent manner, the Hh gradient induces expression of *patched* (*ptc*), *decapentaplegic* (*dpp*), *knot* (*kn*), and *engrailed* (*en*), and regulates the intracellular distribution of *cubitus interruptus* (*ci*) and *smoothened* (*smo*) (Torroja et al., 2005). While signaling in the wing primordium has been extensively studied, little is known about Hh signaling elsewhere in the disc. Hh signaling has not been characterized in the notum primordium or in two other tissues that associate physically and functionally with the notum primordium. These are myoblast progenitors of the adult flight muscles and tracheal tubes that transport air and contain progenitors of adult trachea. The wing disc-associated tracheal progenitors include the air sac primordium (ASP) that will give rise to the adult dorsal air sacs that deliver air to the thoracic flight muscles (Huang and Kornberg, 2015; Sato and Kornberg, 2002).

Both the myoblasts and ASP are situated next to the notum primordium; they are inside the basement membrane that envelopes the disc and are adjacent to the disc epithelium. In this region of the disc, Hh, Dpp, Branchless (Bnl, a FGF), and Wingless (Wg, Wnt homolog) are produced in different, partially overlapping groups of cells (Fig. 1A). Wg signals to myoblasts, and Dpp and Bnl signal to the ASP (Du et al., 2018; Roy et al., 2014). Several targets of Hh signaling are expressed in the notum, but the roles and mechanisms of Hh signaling in the notum and its adjacent cells are unknown.

**Figure 1.**
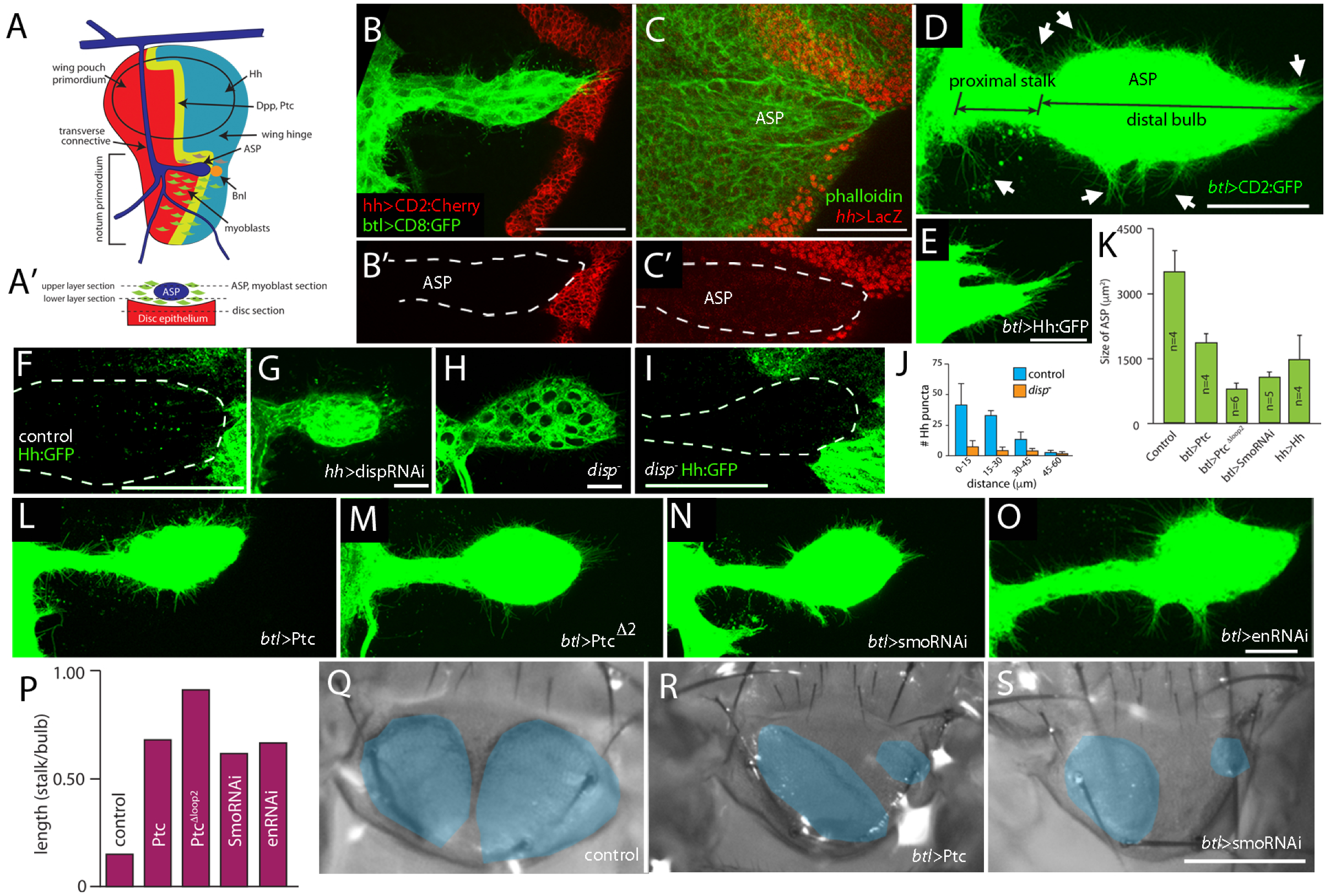
Hh signaling in the ASP. (A) Schematic indicating the relative positions of the wing disc, ASP, and myoblasts, and of *dpp, ptc, bnl*, and *hh* expressing cells; (A’) cross section shows the relative proximity of the ASP, myoblasts and disc epithelium; dashed lines show optical sections that transect the ASP and disc at specified points. (B-C) ASP marked by green fluorescence from *btl-lexA>CD2:GFP* in (B) and Phalloidin in (C), and outlined by white dashed lines in (B’ and C’); *hh* expression marked by red fluorescence in (B; *hh-Gal4>CD8:Cherry*) and in (C; *hh-lacZ*). (B’C’) red channel only; (B,C) merge. Hh expression only detected in the disc (arrows). (D) ASP imaged with green fluorescence (*btl-Gal4>CD8:GFP*) to distinguish the distal bulb, proximal stalk and cytonemes (arrows). (E) Morphologically abnormal ASP with ectopic over-expression of Hh (*btl-Gal4>Hh, mCD8:GFP*). (F) Hh:GFP (green) expressed from BAC in control larva. (J) Graph comparing ASP length in the indicated genotypes. (G) Hh:GFP (green) expressed from BAC in control larva. (H,I) Morphologically abnormal ASPs from a *disp* mutant (H) and larva expressing *dispRNAi* driven by *hh-Gal4*. (J) *disp* mutant ASP (Hh:GFP BAC, *btl>mCD8GFP*) is smaller and has fewer fluorescent puncta than control (F). (J) Decreased number of fluorescent Hh:GFP puncta quantified as a function of proximal/distal position. (n=6, all genotypes). (K-P) ASP morphology is abnormal in genotypes with Ptc (L) and 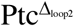 (M), *smoRNAi* (N), and *enRNAi* (O) expression driven by *btl-Gal4* compared to control (D). Degree of size and shape changes are quantified in (K) and (P), respectively. Differences between control and mutants are statistically significant (student’s t-tes P<0.05). (P; n=6, all genotypes). (Q-S) Morphology of adult air sacs (indicated by blue areas) are abnormal in flies over-expressing Ptc (R) or *smoRNAi* (S) driven by *btl>Gal4* in the tracheal system compared to control (Q). Scale bars: (B-C’) 50μm; (D,H,I,L-O) 20μm; (E,G) 50μm; (Q-S) 50μm

Previous studies show that movement of Dpp and Bnl from the disc to the ASP is mediated by cytonemes. Cytonemes are specialized signaling filopodia that transport morphogens and facilitate cell-to-cell transfer of signaling proteins at synapses that link producing and target cells. These synaptic contacts share features with chemical synapses of neurons, including components that localize specifically to pre-synaptic compartments such as the voltage-gated calcium channel, synaptotagmin and synaptobrevin (Huang et al., 2019). Cells that do not make functional synapses do not signal (Huang et al., 2019; Roy et al., 2014). Cytonemes have been observed in many developmental contexts, including Drosophila epithelial, myoblast, and germline cells, between cells in mouse and zebrafish embryos, and vertebrate cells in culture (González-Méndez et al., 2019; Zhang and Scholpp, 2019). In the wing disc, both posterior and anterior compartment cells of the wing pouch project cytonemes that are required for Hh signaling. The ASP projects cytonemes from the tip and medial regions that take up Bnl and Dpp, respectively. Myoblasts project cytonemes that take up Wg produced in the wing disc and project cytonemes that activate Notch signaling in the ASP.

Here, we characterize Hh signaling in the notum primordium and report that Hh from the posterior compartment activates Hh signaling in three target tissues: anterior compartment cells of the notum, the ASP, and the myoblasts. The key findings are that Hh signaling in the ASP is cytoneme-mediated and gene targets of Hh signaling differ in the disc, ASP, and myoblasts.

## Results

### Growth and morphogenesis of the ASP relies on Hh signaling

The ASP is a single cell layered epithelial tube that extends posteriorly across the notum primordium. It develops de novo during the third larval instar from the transverse connective, and its distal tip extends over a small region of the posterior compartment (Fig. 1A-D). Surrogate measures of *hh* gene expression (*hh-lacZ* and *hh-Gal4, UAS-mCherry* transgenes) give no evidence for *hh* transcription in the ASP (Fig. 1B,C), but ectopic over-expression of Hh in the ASP suppressed its growth and morphogenesis (Fig. 1E). This response indicates that the ASP is sensitive to Hh; the closest and most likely source is the disc.

To investigate whether Hh is present in the ASP despite the lack of expression, we monitored ASP preparations using a Hh:GFP BAC transgene that produces fluorescent Hh and is haplosufficient (Chen et al., 2017). We examined normal (control) larvae and detected fluorescent puncta in the ASP, and observed that the amount of Hh:GFP in the ASP is graded, with greater amounts in the cells closest to the Hh-producing disc cells (Fig. 1F). We also examined ASPs isolated from larvae that are deficient for *dispatched (disp)*. Disp is required for Hh release from Hh-producing cells, and because maternal Disp is provided to the embryo, *disp* mutants have sufficient residual function to develop to the L3 stage, albeit with small discs (Burke et al., 1999). Under conditions of *dispRNAi* suppression and in *disp* mutants, growth of the ASP is reduced and ASP morphogenesis is abnormal (Fig. 1G,H). In addition, fluorescence of BAC-encoded Hh:GFP is reduced in the ASPs from *disp* mutant larvae (Fig. 1I,J). These results are consistent with the idea that the ASP takes up Hh from the wing disc and that ASP growth and morphogenesis depends on Hh signaling.

To characterize Hh signaling in the ASP, we genetically modified *ptc, smoothened (smo)*, and *en*, three genes that are Hh targets in other contexts (Fig. 1K-S). Ectopic overexpression of either the normal Ptc protein or a dominant negative mutant 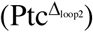 decreases the size of the ASP by 47% and 78%, respectively (Fig. 1K-M). The normal ASP has a short narrow stalk proximal to the transverse connective and a bulbous distal end (Fig. 1 A,B,D). Ectopic expression of Ptc or 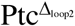 increases the relative length of the stalk by factors of 4.5 and 6, respectively (Fig. 1L,M,P). These results are consistent with the idea that Hh signaling is necessary for normal ASP growth and morphogenesis.

Smo is essential for Hh signaling, and expression of *smoRNAi* decreases Hh signaling in the cells that normally respond to Hh in the wing disc (Casso et al., 2008). To assess the role of Smo in the ASP, we expressed *smoRNAi* in the ASP and observed both that the size of the ASP decreased by 70% and that the relative length of the stalk increased by a factor of 4.1 (Fig.1 K,N). The third known Hh target we monitored is *en*, which encodes a homeodomain-containing transcriptional repressor. Although *en* is a “selector gene” for the posterior compartment (García-Bellido, 1975) where it is a positive regulator of *hh* (Tabata et al., 1992), *en* is also activated by Hh signaling in the wing primordium (Blair, 1992; de Celis and Ruiz-Gómez, 1995; Strigini and Cohen, 1997). We expressed *enRNAi* in the ASP and observed that the size of the ASP decreased by 24% and the relative length of the stalk increased by a factor of 4.4 (Fig. 1K,N,P). These phenotypes are consistent with the idea that *en* is a target of Hh in the ASP and that *en* is necessary for ASP growth and development.

The ASP is the progenitor of the thoracic dorsal air sacs (Sato and Kornberg, 2002), and examination of control WT adult flies detects two almond-shaped dorsal air sac lobes of approximately equal size in the scutellum (Fig. 1Q). In contrast, the dorsal air sacs in flies with Hh signaling suppressed by either ectopic overexpression of Ptc or expression of *smoRNAi* or *enRNAi* are smaller, misshapen, and unequal in size (Fig. 1R,S). In conclusion, phenotypes in conditions of Hh mis-regulation and mutant conditions for *disp, ptc, smo*, and *en* are consistent with the idea that Hh signaling is important to the ASP.

### Graded output of Hh targets in the ASP

We next investigated how genes that are targets of Hh signaling in other tissues are expressed in the ASP, and examined mutant contexts to determine if the expression patterns of the targets are consistent with the idea that their expression in the ASP is regulated by Hh. We examined Ptc, Smo, En, and Cubitus interruptus (Ci, the transcription factor that mediates transcriptional outputs of Hh signal transduction), and the Hh co-receptor Interference hedgehog (Ihog) (Lum et al., 2003). In other contexts, Hh signaling elevates Ptc, Smo and En amounts, and suppresses Ci and Ihog.

To determine if *ptc* expression changes in response to different levels of Hh signaling, we monitored Ptc protein by fluorescence of BAC-encoded Ptc:Cherry and *ptc* transcription by fluorescence of nuclear GFP expressed under the control of a segment of the *ptc* regulatory region. These assays are quantitative and reveal that Ptc protein and *ptc* transcription are elevated in cells at the distal tip of the ASP. These cells are the closest to Hh-expressing disc cells. Proximal ASP cells, which are more distant, express lower levels (Fig. 2A,B). To examine Smo, we monitored fluorescence of BAC-encoded Smo:GFP (Chen et al., 2017). In the anterior compartment of the wing pouch primordium, *smo* expression is uniform but amounts of Smo protein are higher in cells that activate Hh transduction (Alcedo et al., 2000), and as shown in Figure 2C, the distal ASP cells have higher amounts of Smo. To determine the distribution of En in the ASP, we applied α-En antibody and observed staining that is strongest in the distal bulb and absent in the proximal stalk (Fig. 1D and 2C). In contrast, fluorescence of Cherry:Ci (Fig. S1) has the opposite contour (lower in distal cells and elevated in proximal cells; Fig. 2E) and α-Ihog antibody staining is constant across the ASP proximal-distal axis (Fig. 2F).

**Figure 2.**
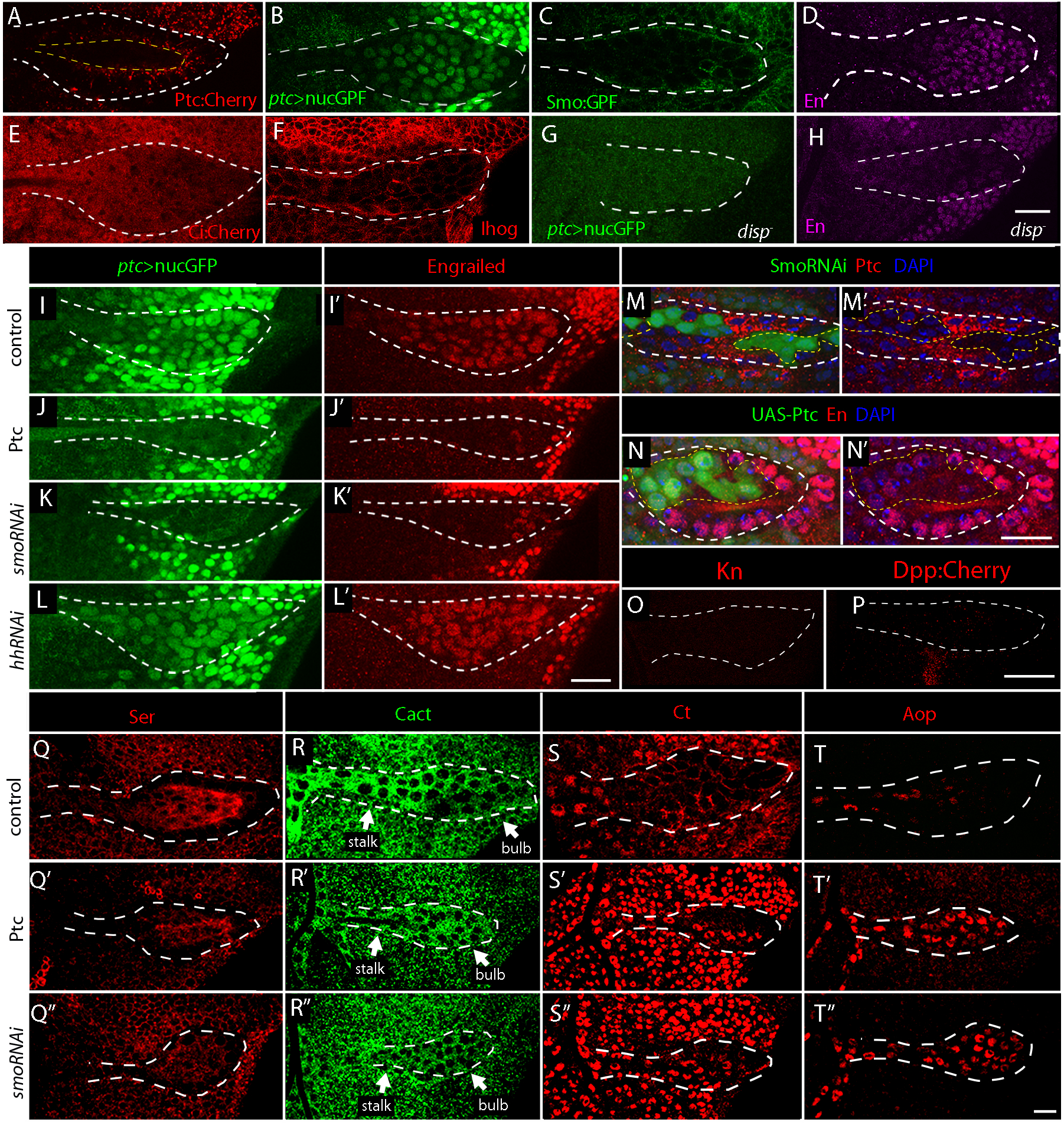
Graded output of Hh targets in the ASP. (A-F) Distributions of (A) Ptc:Cherry, (B) ptc:nucGFP, (C) Smo:GFP BAC, (D) α-En, (E) Ci:Cherry Crispr, (F) Ihog:Cherry in the ASP. Majority of the Ptc:Cherry localizes to the apical lumen of the ASP (A, yellow dashed lines). In all panels, the ASP is outline with white dashed lines.(G and H) Intensity of ptc:nucGFP fluorescence (G) and anti-En staining (H) is decreased in *disp*^*-*^ mutants compared to control in (B) and (D), respectively. (I-L) Fluorescence of ptc>nucGFP (green) is decreased upon the overexpression of *Ptc* or *smoRNAi* in the tracheal system (J: *btl>Ptc* and K: *btl>smoRNAi*, respectively) compared to control (I) or upon *hhRNAi* expression of in the trachea (*btl-Gal4>hhRNAi*, L). (I’-L’) Intensity of anti-En (red) staining decreases in the ASP when *Ptc* or *smoRNAi* are overexpressed in the tracheal system (J’-*btl>Ptc* and K’-*btl>smoRNAi*, respectively) compared to control (I’) or when *hhRNAi* is expressed in the tracheal system (*btl-Gal4>hhRNAi*, L’). (M and M’) In the ASP, outtlined with white dashed line, levels of anti-Ptc staining (red) is decreases in clonal cells expressing *smoRNAi* (green) compared to its surrounding control cells (without green). (N and N’) In the ASP, white dashed line, levels of anti-En staining (red) is decreases in clonal cells expressing *Ptc* (green) compared to its surrounding control cells (without green). (O-P) Signals of α-Kn antibody staining and Dpp:Cherry is absent in the ASP (outlined by white dashed line). (-Q-Q’’) High levels of anti-Ser (red) staining is observed in the bulb of the ASP. The levels of anti-Ser is lower in the ASP of larvae that express *Ptc* or *smoRNAi* in the trachea system (*btl-Gal4>*Ptc-S’ and *btl>smoRNAi*-S’’). (R-R’’) Fluorescence intensity of Cact:GFP (green) is stronger in the stalk and TC compared to the bulb of the ASP in control larvae (R). In larvae over-expressing Ptc (R’, *btl-Gal4>*Ptc) or *smoRNAi* (*btl>smoRNAi*, R’’) in the tracheal system, the fluorescence pattern of Cact:GFP became uniform in the ASP. (S-S’’) Nuclear staining of α-Ct is seen in the stalk and the TC but absent in the bulb in the ASP of control larvae. (S). In larvae over-expressing Ptc (*btl-Gal4>*Ptc, S’) or *smoRNAi* (*btl>smoRNAi*, S’’) in the tracheal system, the nuclear Ct can be seen in both in the bulb and stalk. (T-T’’) Nuclear staining of α-Aop is seen in the stalk and the TC but absent in the bulb in the ASP of control larvae(T). In larvae over-expressing Ptc (*btl-Gal4>*Ptc, T’) or *smoRNAi* (*btl>smoRNAi*, T’’) in the tracheal system, the nuclear Ct can be seen in both in the bulb and stalk. In all panels, white dashed line outlines the ASP. Scale bars: (H,L’,N’,P) 20μm; (T’’) 50μm.

To determine if expression of *ptc* and *en* in the ASP are dependent on Hh signaling, we monitored mutants for *ptc* and En expression (Fig. 2-N’). In *disp* mutants, both the fluorescence of the *ptc*-GFP reporter and the intensity of En staining is reduced in the ASP (Fig. 2G,H). Ectopic overexpression of Ptc or *smoRNAi* in the ASP with the *breathless* (*btl)* pan-tracheal driver (Fig. 2I-L’) or in somatic clones (Fig. 2M-N’), also reduces *ptc* expression and En. In contrast, ectopic over-expression of *hhRNAi* in the ASP (which does not express *hh*), does not change expression of the *ptc* reporter or En (Fig. 2L,L’). Kn and Dpp are two additional targets of Hh signaling in the wing pouch. In the ASP, we did not detect signals of α-Kn antibody staining and fluorescence of Dpp:Cherry Crispr knock-in, suggesting that they are not targets in this tissue (Fig. 2O and P). We also observed that amounts of Serrate (Ser), Cactus (Cact), Cut (ct), and Anterior open (aop)are elevated in the more proximal ASP cells relative to distal ASP cells, and that the low amounts in the more distal ASP cells are sensitive to Hh signaling (Fig. 2Q-T”). We conclude that Hh functions as a paracrine signaling protein that generates graded distributions of Ptc, Smo, En, Ser, Cact, and Aop targets in the ASP.

### Hh produced in the wing disc induces signal transduction in myoblasts

During the third instar period, myoblasts proliferate and spread across the wing disc to cover approximately 2/3 of the notum primodium (Fig. 1A) (Gunage et al., 2014; Huang and Kornberg, 2015). Approximately 125 myoblasts lie directly over Hh-expressing, posterior compartment disc cells near the wing hinge primordium (Fig. 3A). This region is also near the distal tip of the ASP. We investigated whether these myoblasts express Hh or might respond to Hh made by the wing disc. We examined the expression of two surrogate sensors for *hh* expression, *hh-lacZ* and *hh-Gal4*, and did not detect activity (Fig. 3B-C’). We did, however, find evidence for Hh signaling. We detected *disp*-dependent fluorescence of Hh:GFP or Smo:GFP in myoblasts that are near the posterior compartment in animals that express either of these proteins from BAC transgenes (Chen et al., 2017) (Fig. 3D-E’’), and in preparations probed with antibodies directed against either Ptc or En (Fig. 3F-G’). The amounts of Hh, Smo, Ptc, and En in the myoblasts vary with position: they are greatest in myoblasts closest to the Hh expressing cells of the wing disc posterior compartment and are lower in myoblasts that are more distant and overlie the nearby anterior compartment. We also tested for expression of Ct, Aop, Cact, Ser, Dpp, and Kn, which are Hh targets in other tissues. Ct, Cact, and Aop are expressed uniformly in the myoblasts and do not appear to be Hh targets (Fig. 3H-J). We did not detect expression of either Ser, Dpp, or Kn (Fig. 3K-M). These findings are consistent with the idea that the myoblasts that in the immediate vicinity of the posterior compartment take up Hh and activate Hh signal transduction.

**Figure 3.**
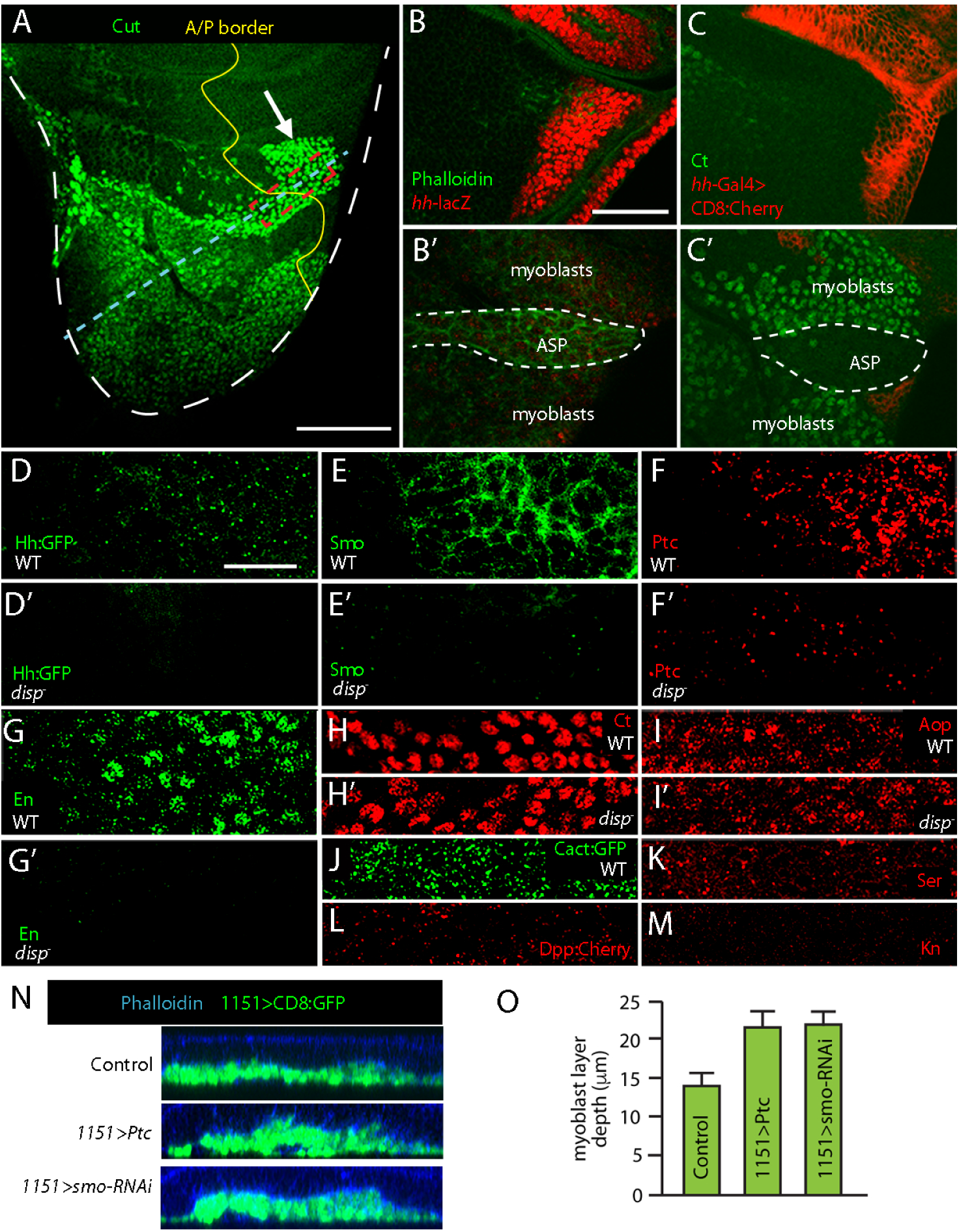
Wing disc derived Hh activates signal transduction in the myoblasts. (A) The myoblasts (marked by α-Ct staining, green) populate the notum region of the wing disc (white dashed line) in late L3 larvae. A population of myoblasts (arrow) resides very close to the *hh* expressing cells of the wing disc (right side of yellow dashed line). (B-C’) Expression of *hh-lacZ* (red in B) and *hh-Gal4* (red in C) can be seen in the notum. However, the expression of *hh-lacZ* and *hh-Gal4* cannot be seen in either the myoblast or the ASP. (D-M) Distribution of Hh targets within the myoblasts in the dashed red lined box in (A). (D-G’) Distributions of Hh:GFP BAC fluorescence (D), α-Ptc staining (E), Smo:GFP BAC fluorescence (F), and α-En-staining (G) in myoblasts of wild-type and in disp mutant larvae (D’, E’, F’, G’ respectively.) Levels of signals for all four are decreased in *disp*^-^ mutants compared to control. (H-M) Distribution of α-Ct staining (H), and α-Aop staining (I) in wild-type. These levels do not change in *disp*^*-*^ mutants (H’ and I’respectively). (J-M) Cac:GFP fluorescence, α-Ser staining, Dpp:Cherry fluorescence, α-Kn staining shows uniform distribution. (N) Orthogonal sections of control (*1151-gal4>+*), *1151>Ptc* (*1151-Gal4, UAS-Ptc*), and *1151>smoRNAi* (*1151Gal4, UAS-smoRNAi*), wing discs and myoblasts along the blue dashed line shown in panel A. The outline of the disc is shown with phalloidin staining (blue) and the layers of myoblasts are visualized with the expression of CD8:GFP in the myoblasts (*1151-Gal4, UAS-CD8:GFP*). (O) Graphs showing the depth of the myoblast layers for genotypes that knockdown Hh signaling. Difference between control and *1151>Ptc*, and *1151>smoRNAi* are statistically significant (student’s t-test, P value<0.05); n=5. Scale bars: (A) 100μm; (B) 50μm; (D) 20μm.

To investigate whether Hh signaling has a role in myoblast development, we analyzed the myoblasts in genetic conditions that disrupt Hh signaling. Although the myoblasts constitute several cell layers that form a stack of approximately 14 μm at late third instar (Gunage et al., 2014), we found that the depth of this myoblast layer increased under conditions of myoblast overexpression of Ptc or *smoRNAi* (Fig. 3N-O). This phenotype is consistent with the idea that Hh signaling has a formative role for the myoblasts.

### Hh signaling in the notum primordium

To investigate Hh signaling in the notum primordium, we monitored the expression of genes which are Hh targets in other tissues. Ptc is a target of Hh signaling in the notum primordium, as it is in the wing pouch and all other known contexts. *en* is another target. Its expression in the anterior compartment of the wing pouch primordium is activated by Hh signaling (de Celis and Ruiz-Gómez, 1995; Strigini and Cohen, 1997), and as shown in Figure 2, Hh signaling activates *en* expression in the ASP (Fig. 2D,H,I-L’). We stained third instar wing discs with α-En antibody and observed that En is present near the distal tip of the ASP both in the posterior compartment as well as in approximately 175 (n=4, st.dv.=21) cells on the anterior side of the compartment boundary (Fig. 4A,B). To test if En expression in the anterior compartment is Hh-dependent, we examined *disp* mutants and detected significantly lower amounts of En than in controls (Fig. 4B,C). Additional evidence for Hh signaling in this region was obtained by monitoring Ci. Fluorescence of Cherry:Ci in anterior compartment cells that express En is reduced relative to more anterior portions of the disc (Fig. 4A’). This reduction of Ci amounts is *disp*-dependent (Fig. 4B’-C”), consistent with the idea that Hh signaling regulates both En and Ci in these cells. We also analyzed expression of Dpp in the notum primordium and tested if it is under the control of Hh. Dpp is expressed in a stripe of cells, with the majority overlapping with or alongside cells that express Ptc. Expression of Dpp is significantly lower in *disp* mutants, consistent with the idea that Dpp expression is dependent on Hh signaling. In contrast, Ser, Cact, Ct, and Aop, which are targets of Hh signaling in the ASP (Fig. 2Q-T) are not expressed in the notum primordium (Fig. 3), suggesting that they are not targets of Hh signaling in the notum.

**Figure 4.**
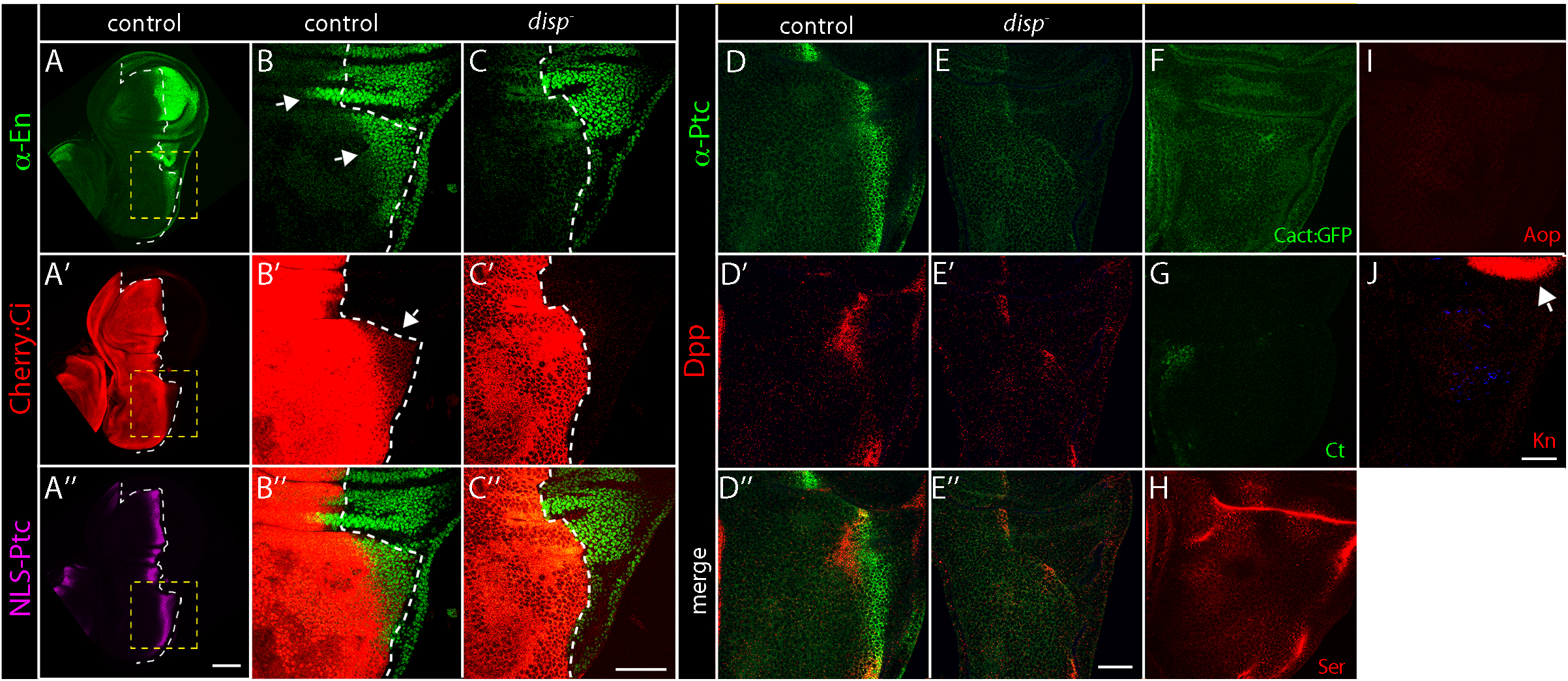
Targets of Hh signaling in the notum. (A-A’’) Distributions of α-En staining (A), Cherry:Ci CRISPR fluorescence (A’), nuclear>Ptc fluorescence (A’’) in the wing disc. The compartment boundary is marked by the white dashed line. (B-B’’) Anti-En (green) and Cherry:Ci fluorescence (red) in the notum portion of the wing disc that is marked by a yellow dashed line in (A). The compartment boundary is marked by white dashed line. Anti-En staining can be detected in the anterior compartment (arrows in B) and Cherry:Ci fluorescence staining is decreased in the anterior compartment cells that are adjacent to the compartment boundary. (C-C’’) α-En staining in the anterior compartment and the decrease of Cherry:Ci fluorescence at the compartment boundary is diminished in *disp*^-^ mutants. (D-E’’) α-Ptc staining (green) and α-Dpp staining (red) in wild-type or in *disp* mutant notum primordium. Arrows point to staining in the notum primordium. (F-J) Notum primordium showing the fluorescence of Cact:GFP, and α-Ct, α-Ser, α-Aop, and α-Kn staining. Scale bars: (A’’) 100μm; all others: 50μm.

### Hh signaling in the ASP is mediated by cytonemes

To investigate whether uptake of Hh by ASP cells is cytoneme-mediated, we imaged preparations from animals that express membrane-tethered fluorescent proteins. Figure 5A,B shows that cytonemes containing CD4:mIFP (Yu et al., 2015) extend from the ASP tip and take up Hh:GFP from the wing disc. Cytonemes also extend from the medial region of the ASP, but these cytonemes do not extend toward Hh-expressing disc cells and do not take up Hh:GFP. Time-lapse imaging shows that cytonemes that extend from the ASP tip contain motile puncta that have Hh:GFP (Fig. 5C) and Ptc:Cherry (Fig. 5D). These results are consistent with the possibility that ASP cytonemes transport Hh from the disc to the ASP.

**Figure 5.**
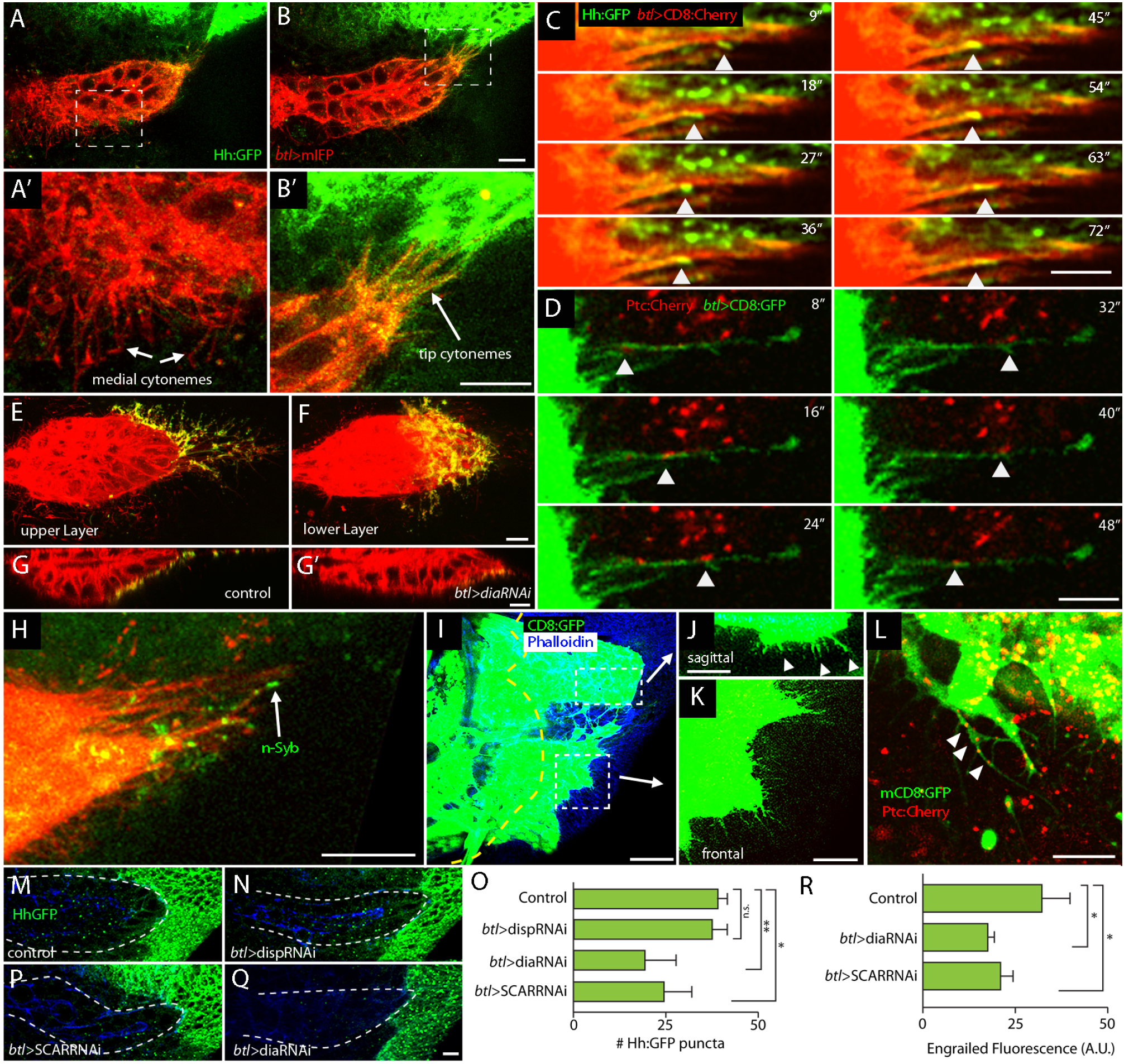
Cytonemes mediate Hh signaling between the ASP, myoblast and the notum. (A-B’) ASP of third instar larvae expressing *mIFP* in the ASP (red, *btl>mIFP*) and *Hh:GFP BAC* (green). A and B shows optical section with medical cytoneme (A) and tip cytoneme (B). (A’ and B’) High magnifications of the boxed regions in A and B show strong accumulation of Hh:GFP in the tip cytonemes but not in the medial cytonemes. (C) Different time points of time-lapse videos shows Hh:GFP puncta (arrow) moving along ASP cytoneme (red, marked by *btl>CD8:Cherry*). (D) Snapshots of time-lapse video shows Ptc:Cherry fluorescent puncta (arrow) moving along ASP cytoneme (green, marked by *btl>CD8:GFP*). (E-F) GRASP signal (green) shows contact between the ASP cytonemes (red) and Hh producing cells in both the upper and lower layer of the ASP. Genotype: *btl>Cherry:CAAX, CD4:GFP*^*1-10*^, *hh>CD4:GFP*^*11*^. (G-G’) Sagittal view of ASP shows that the number of cell-cell contacts (green) decreases upon expression of *diaRNAi* (G’) in the tracheal system compared to control (G). Genotypes: (G)-*btl>Cherry:CAAX, CD4:GFP*^*1-10*^, *hh>CD4:GFP*^*11*^, (G’)-*btl>Cherry:CAAX, CD4:GFP*^*1-10*^, *diaRNAi, hh>GFP*^*11*^. (H) n-syb GRASP (green) show that cytonemes form active synapses with the *hh* producing cells of the wing disc. Genotype: *btl>Cherry:CAAX, CD4:GFP*^*1-10*^, *hh>n-syb:GFP*^*11*^. (I-K) Myoblasts (green, *1151-Gal4>mCD8:GFP*) populate the notum region of the wing disc (blue, phalloidin). Yellow dashed line marks the compartment boundary. Cytonemes from the myoblasts can be seen projecting toward the notum posterior compartment in magnified sagittal sections (J, top dasheded white line box) and (K, bottom dashed white lined box). (L) Ptc:Cherry puncta localize to myoblast cytonemes (arrows). Genotype: *1151-Gal4>mCD8:GFP, Ptc:Cherry BAC*. (M-Q) Compared to control (*btl>+, Gal80*^*ts*^) and to *disp* reduction (*btl>dispRNAi, Gal80*^*ts*^), the number of Hh:GFP puncta (green) is decreased upon downregulation of *scar* or *dia* (*btl>scarRNAi, Gal80*^*ts*^ and *btl>diaRNAi, Gal80*^*ts*^). (O) Graph showing number of Hh:GFP puncta in control (*btl>dispRNAi, Gal80*^*ts*^, *btl>diaRNAi, Gal80*^*ts*^), and *btl>ScarRNAi, Gal80*^*ts*^. n=4-6 (R) Graph showing levels of α-En staining in ASP in control (*btl>+, Gal80*^*ts*^), or downregulation of *dia* or *scar* (*btl>scarRNAi, Gal80*^*ts*^ and *btl>diaRNAi, Gal80*^*ts*^). α-En staining decreases upon *dia* or *scar* knock-down. Student’s t-test P values: *<0.05, ** <0.005, n.s. not significant. n=5. All scale bars: 10μm, except (I) 50μm.

GRASP (GFP reconstitution across synaptic partners) is a technique that generates fluorescence at contacts made by cells that express membrane-tethered, extracellular fragments of GFP (transmembrane domain of mouse CD4 fused to GFP^1-10^ and to GFP^11^) that together can reconstitute fluorescent GFP functional photocenter if they are stably and closely juxtaposed (Feinberg et al., 2008). GRASP fluorescence marks contacts between disc cells that express Dpp and disc cells that take up Dpp (Roy et al., 2014), between disc cells that express Hh and disc cells that take up Hh (Chen et al., 2017; González-Méndez et al., 2017), and between disc cells that make Bnl and ASP cells that take up Bnl (Roy et al., 2014). To test whether ASP cytonemes contact disc cells that produce Hh, we expressed CD4:GFP^1-10^ in the ASP and CD4:GFP^11^ in Hh-expressing disc cells, and marked ASP cytonemes with membrane-tethered Cherry (Cherry:CAAX). In optical sections that image the ASP at distances farthest from the basal surface of the ASP cells (“upper layer”; Fig. 1A’), we observed GFP fluorescence along ASP cytonemes (Fig. 5E). In optical sections at the lower layer of the ASP or in sagittal sections, GRASP fluorescence is present in the region between the ASP and wing disc (Fig. 5F,G). We also monitored the GRASP-marked contacts in genetic conditions that compromise cytonemes. Pulsed expression of RNAi directed against *diaphanous* (*dia*), a formin-encoding gene, decreases the number of cytonemes without affecting cell viability or cell polarity (Bischoff et al., 2013; Chen et al., 2017; Roy et al., 2014); and it reduces fluorescence of CD4:GFP^11^ – CD4:GFP^1-10^ GRASP (Fig. 5G,G’). Further evidence for the nature of the contacts that ASP cytonemes make with Hh-producing disc cells was obtained by using a variant GRASP system that substitutes the cytoplasmic and transmembrane domains of n-Synaptobrevin (Syb), a constituent of synaptic vesicles, for the CD4 portion of CD4:GFP^1-10^. Syb GRASP identifies active synapses in neuronal tissue (Macpherson et al., 2015), and marks contacts between Hh-producing and Hh-receiving cells in the wing disc (Chen et al., 2017). As shown in Figure 5H, expression of Syb:GFP^1-10^ in wing disc posterior compartment and expression of CD4:GFP^11^ in the ASP generates fluorescence at contacts between ASP cytonemes and Hh-expressing disc cells. In sum, these experiments show that cytonemes at the ASP tip synapse with Hh-expressing disc cells and are consistent with the idea that Hh moves from the disc to the ASP at these contacts.

The myoblasts also extend cytonemes to the disc (Fig. 5I-L). Some orient directly toward underlying disc cells and are visible in sagittal sections (Fig. 5J); others extend across the disc and are visible in frontal sections (Fig. 5K). Punctal Cherry fluorescence is visible in myoblast cytonemes in animals that have the Ptc:Cherry-encoding BAC (Fig. 5L).

To obtain evidence for cytoneme-mediated Hh uptake, we expressed RNAi transgenes that decrease *disp, dia*, and *SCAR* expression, and monitored both the presence of Hh and Hh signaling in the ASP. Previous studies establish that these conditions of *diaRNAi* and *SCARRNAi* expression decrease cytoneme-mediated signaling without disrupting cell polarity, morphology, or viability (Bischoff et al., 2013; Chen et al., 2017; Roy et al., 2014). In controls and in animals that express *dispRNAi* in the ASP, Hh:GFP fluorescence is present in similar amounts (Fig. 5M-O). In contrast, Hh:GFP fluorescence in the ASP is reduced by *disp* knockdown in the disc posterior compartment (Fig. 1I). Knockdown of *dia* or *SCAR* in the ASP decreases Hh:GFP fluorescence by approximately 50% (Fig. 5O-Q), consistent with the idea that cytoneme ablation decreases transport of Hh to the ASP. We monitored Hh signal transduction by staining for En, and observed that knockdown of either *dia* or *SCAR* in the ASP decreases amounts of En in the ASP by >50% (Fig. 5R). These results are consistent with the idea that cytonemes are required for Hh transport to the ASP and for Hh pathway activation in the ASP.

ASP cytonemes that take up Hh from the disc extend from the distal tip (Fig. 5). Because previous studies showed that Bnl is also taken up from the disc by tip cytonemes, in contrast to the cytonemes that extend from medial regions and take up Dpp (Roy et al., 2014), we investigated cytoneme-mediated uptake of Hh in more detail. We first examined preparations from flies the express BAC-encoded Hh:GFP (Chen et al., 2017) and Btl:Cherry generated by CRISPR-generated knock-in (Du et al., 2018). Both Bnl-expressing cells and Hh-expressing cells are distal to the ASP tip, and we observed that ASP tip cytonemes in these preparations contain both Hh:GFP and Btl:Cherry (Fig. 6A). This indicates that tip cytonemes might take up Hh and Bnl from the disc and might have receptors for Hh and Bnl. We examined preparations from animals with *btl*-Gal4 driven over-expression of Ptc:GFP and Btl:Cherry, and detected both Ptc:GFP and Btl:Cherry in tip cytonemes (Fig. 6B). These results are consistent with the idea that tip cytonemes might take up either or both signaling proteins.

**Figure 6.**
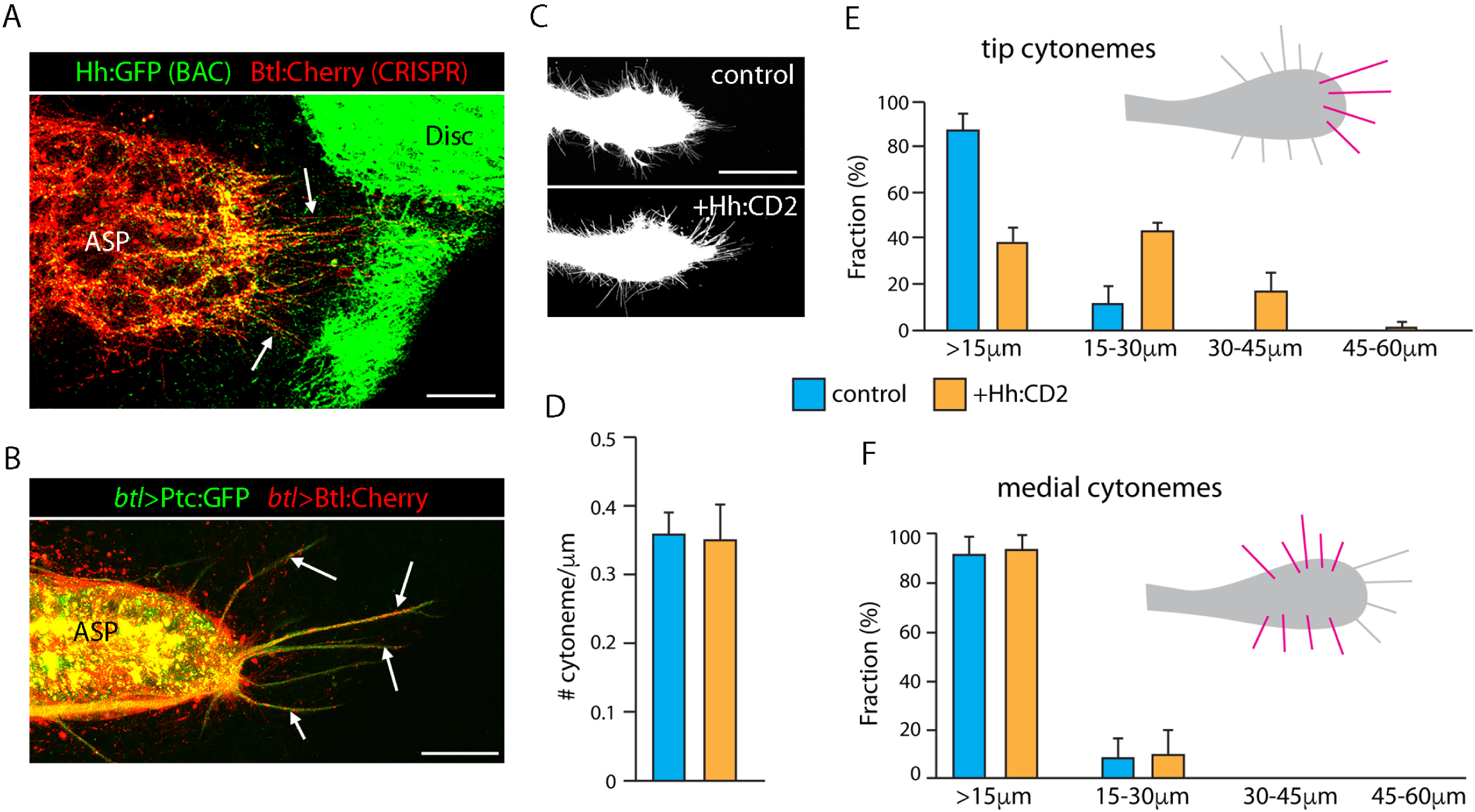
Cytonemes respond to Hh signaling. (A) Image of an ASP and wing disc from larva that expresses BAC encoded Hh:GFP (green) and CRISPR-generated Btl:Cherry knock-in (red). Arrows point to tip cytonemes marked by Btl:Cherry and Hh:GFP. Genotype: *Hh:GFP*/+; *Btl:Cherry*/+. Scale bar: 25μm. (B) Image of an ASP that expresses Ptc:GFP (green) and Btl:Cherry (red) driven by *btl-Gal4*. Arrows point to tip cytonemes that contain both Ptc:GFP and Btl:Cherry. Genotype: *btl-Gal4, UAS-Btl:Cherry; UAS-Ptc:GFP/+.* (C) Images of ASPs from larvae with *btl-Gal4* and *UAS-mCD8:GFP* only (control) or together with *UAS-Hh:CD2*. Genotypes: *btl-Gal4, UAS-mCD8:GFP/+; Gal80ts/+* and *btl-Gal4, UAS-mCD8:GFP/+; Gal80ts/UAS-Hh:CD2.* Scale bar: 50 μm. (D-F) Graphs showing (D) density of ASP cytonemes measured as # cytonemes/μm of ASP perimeter, and fraction of cytonemes of indicated lengths at the ASP tip (E) and ASP medial region (F). Difference between Control and *UAS-Hh:CD2* in (D) is not statistically significant (student’s t-test P>0.05). Differences between Control and *UAS-Hh:CD2* are statistically significant for tip cytonemes (E; student’s t-test P<0.005) but not for medial cytoenemes (F; student’s t-test P>0.05). n=4. Scale bar: 25μm.

We next analyzed cytonemes extending from ASPs that ectopically over-express Hh:CD2, an engineered Hh that substitutes the transmembrane domain of the mouse CD2 protein for the C-terminal cholesterol of mature Hh (Strigini and Cohen, 1997). This membrane-tethered, chimeric protein has signaling activity, but unlike the normal protein does not move efficiently beyond the cells that express it. Because the ASP does not grow normally in the presence of constitutive ectopic over-expression of Hh:GFP (Fig. 1E) or Hh:CD2 (not shown), we expressed Hh:CD2 for a limited time during the third instar by conditional inactivation of Gal80^ts^. Under this regimen of Hh:CD2 expression, the ASP grows to normal size and morphology to late third instar at permissive temperature (18°C), and cytonemes are analyzed after 24 hours at non-permissive temperature (29°C). We found that ectopic over-expression of Hh:CD2 did not change either the total number of ASP cytonemes or the lengths of medial cytonemes relative to controls. However, the average length of tip cytonemes increased (Fig. 6C-F). In this experiment, because Hh:CD2 was expressed only in tracheal cells and the disc was genetically unaltered, this significant cytoneme phenotype is evidence that Hh signaling in the ASP cells that take up Hh from disc contributes to cytoneme production or behavior. This effect is consistent with the conclusion that the tip cytonemes mediate Hh uptake from the disc and with the positive feedback model of Du et al, 2018 that links cytoneme production and stability to signaling.

### Hh signals to disc, ASP and myoblast cells

The findings presented here show that Hh produced in the disc signals to the ASP (Fig. 1) and myoblasts (Fig. 3), and that both types of responding cells are near the wing hinge region of the disc in close proximity to disc posterior compartment cells of the notum primordium (Fig. 1A). The cells in this region therefore present a setting in which a single source of Hh activates signal transduction in three different types of cells – wing disc anterior compartment cells, ASP cells, and myoblasts (Fig. 7A). The confocal microscope-acquired images in the three panels of Figure 7B show the Hh-responding (*ptc*-expressing) cells in two optical sections, one that focuses on the disc epithelium (Fig. 7B) and another that focuses on the plane that includes ASP and myoblast cells (Fig. 7B’). The anterior extent of Hh signaling in the disc is not constant in this region, but the anterior extent of signaling is similar in the disc and ASP where they are juxtaposed, and is similar in the disc and myoblasts where they are juxtaposed. Myoblasts that overlie posterior compartment disc cells also activate Hh signal transduction, although the disc cells block autocrine Hh signaling (Ramírez-Weber et al., 2000).

**Figure 7.**
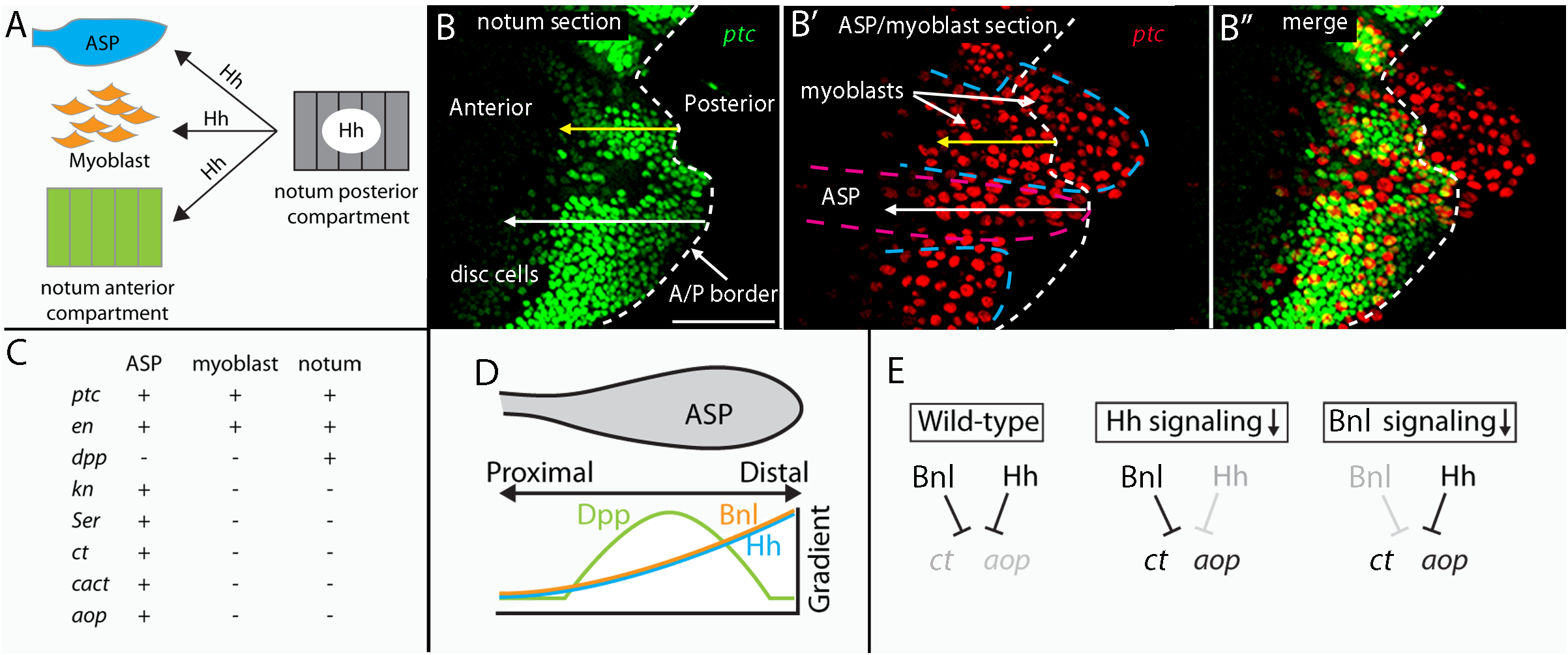
Hh signals to multiple targets. (A) Hh produced by the disc signals to disc cells in the notum, myoblasts and the ASP (B) *ptc*-expressing cells in the disc, ASP, and myoblast, detected by the fluorescence of *ptc*-nuclear GFP in two optical sections. GFP fluorescence is green in the notum section and is red in the ASP/myoblast section. The anterior/posterior compartment border is indicated by dashed white line and the extent of ptc expression in the anterior compartment by the white and yellow arrows. Scale bar: 25μm. (C) Transcriptional targets of Hh signaling in the myoblast, ASP, and notum. (D) Cartoon representing the distributions of Dpp, Bnl, and Hh taken up in the ASP. (E) Cartoon depicting the roles of Bnl and Hh regulating expression of *ct* and *aop* in the ASP of wild type and mutant genotypes that decrease Hh or Bnl signaling.

Another difference in Hh’s action on the three cell types is the suite of target genes that Hh signal transduction regulates (Fig. 7C). Whereas all express *ptc* and *en*, only disc cells express *dpp* and only ASP cells regulate *kn, Ser, ct, cact*, and *aop* in response to Hh (Figs. 1-3). *ct* and *aop* expression in the ASP is also regulated by Bnl signaling (Du et al., 2018). Previous studies identified concentration gradients of Dpp and Bnl that have distinct profiles in the ASP (Du et al., 2018; Roy et al., 2014). The amount of Dpp is highest in the medial region and declines bidirectionally along the proximal-distal axis, but Bnl is highest distally and declines proximally (Fig. 7D). The Hh concentration profile in the ASP (Fig. 1J) has a similar contour to that of Bnl (Fig. 7D), and both Hh signal transduction and Bnl signal transduction are required to regulate *ct* and *aop* (Fig. 7E).

## Discussion

We show in this study that Hh signaling is activated in three tissues: the tracheal ASP and wing disc-associated myoblasts in addition to anterior compartment cells of the notum (Fig. 7). These findings are not unexpected as most Drosophila organs rely on Hh signaling. There are, nevertheless, notable aspects of Hh signaling in these particular tissues that the anatomy of the system reveals. During the 3^rd^ instar period as the ASP grows across the basal surface of the disc toward the posterior compartment where Hh is expressed, the growth and morphology of the ASP is dependent on Hh it takes up from the disc. During this period, myoblasts also proliferate and migrate over the disc basal surface, and they spread around and under the ASP, and myoblasts that are in the region where Hh moves from the disc to the ASP depend on Hh they take up to regulate their growth. And anterior compartment wing disc cells express Dpp in response to Hh they receive, establishing the functionality of a signaling center along the compartment border. Because the wing disc and its associated trachea and myoblasts are physically separate from other tissues inside the larva, and the only local Hh-expressing cells are the cells of the disc posterior compartment, we assume that the posterior compartment disc cells are the relevant source and that Hh from these cells in the wing disc activates signal transduction in these three discreet tissues. Hh produced in the disc is released basally from the polarized columnar epithelial cells (Callejo et al., 2011), and it has been shown to be taken up basally by wing disc cells (Callejo et al., 2011; Chen et al., 2017; González-Méndez et al., 2017). The basal location of the ASP and myoblasts is consistent with the idea that they take up Hh from the same source.

Whereas the number of different morphogen proteins is small - in Drosophila it may be only the four signaling proteins Decapentaplegic (a BMP family member), Wingless, Fibroblast growth factor, and Epidermal growth factor in addition to Hh – the number of tissues that they control and the number of different morphologies they mold is many times larger. The apparent versatility of this system of growth and morphogenesis is presumably a function of the context dependence of signaling, and the three target tissues we analyzed that are activated by disc-produced Hh each express a different suite of target genes(Fig. 7C). Several are targets in only one tissue (e.g., Dpp in the wing; Ser, Cact, Ct, and Aop in the ASP), and only two are targets in all three (e.g., Ptc and En) (Figs. 2,3,4,6,7). It is worth noting that the distance over which Hh signals is also similar in these tissues. There is almost complete spatial overlap between Ptc-expressing cells of the ASP and Ptc-expressing cells of the disc anterior compartment that they overlay, and there is almost complete spatial overlap between myoblasts that overlie the disc anterior compartment and the underlying disc cells (Fig. 7B). We also note that whereas disc cells in the posterior compartment do not activate Hh signal transduction, the myoblasts that overlie Hh-expressing posterior compartment cells do (Fig. 7B). We consider this in the broader context of the tissue specific responses we found, and suggest that they are not likely due to the properties of Hh gradients themselves. Rather, the different target signatures of each tissue may be manifestations of the distinctiveness of each cell type, and in this instance to the *engrailed*-dependent inhibition of autocrine Hh signaling in the disc posterior compartment (Ramírez-Weber et al., 2000).

The band of cells that activates Hh signaling in the disc is not uniformly wide, but is smaller in some regions than others (see for example Figure 1E in (Chen et al., 2017)). A portion of the band is particularly small near the ASP and Hh-responsive myoblasts (Fig. 3), and these differences in the widths of the band of disc cells that activate Hh signaling are also reflected in the extent of Hh signaling in the ASP and myoblasts. We do not understand the basis for these variations and are not aware of physical or morphological features that might differentially affect Hh dispersion in these regions. Hh signaling is cytoneme-mediated (Bischoff et al., 2013; Chen et al., 2017; González-Méndez et al., 2019), and several studies have shown that cytoneme distributions across tissues parallel signaling. In the wing disc and abdominal histoblasts, the lengths and numbers of cytonemes correlates with graded responses to Hh signaling (Bischoff et al., 2013; Chen et al., 2017; González-Méndez et al., 2017), and cytoneme distributions and Bnl signaling are similarly correlated in the ASP (Du et al., 2018). The model proposed by Du et al, 2018 accounts for the matching gradients of cytonemes, morphogen protein amounts, and signaling with evidence that “morphogens self-generate precise tissue-specific gradient contours through feedback regulation of cytoneme-mediated dispersion”. This model invokes an essential role by the target cell in cytoneme propagation and/or stability, a functionality that involves positive-feedback to signal transduction and is supported by our observation that activation of Hh signaling by ectopic expression of Hh:CD2 in the ASP increases the lengths of ASP cytonemes (Fig. 6E).

Although multiple signaling pathways have been shown to be important in tissue regeneration and immune responses (Houtz et al., 2017; Jiang et al., 2009) and in Drosophila leg and eye development (Estella et al., 2008; Newcomb et al., 2018; Tomlinson and Struhl, 2001), we have little understanding how multiple signaling inputs might be coordinated for an integrated output. The ASP receives Dpp, Bnl, and Hh from the wing disc, and all distribute spatially in the ASP in concentration gradients. Whereas highest levels of the Dpp gradient are in the medial region of the ASP (Roy et al., 2011), the Bnl and Hh gradients both extend along the proximo-distal axis with highest levels at the tip of the ASP ((Du et al., 2018) and Fig. 7D). These distributions imply that ASP cells receive similarly graded inputs of both Bnl and Hh, which is consistent with our finding that some tip cytonemes take up both Bnl and Hh (Fig. 6A,B). We found that *ct* and *aop*, which are both targets of Bnl and Hh signal transduction, have similar patterns of expression along the proximo-distal axis. *ct* and *aop* are normally expressed in the proximal stalk region and not in the more distal bulb. However, in ASPs with reduced Bnl or Hh signaling, Ct and Aop expression are elevated in the bulb, thus revealing that suppression of Ct and Aop expression in the bulb is dependent on both Bnl and Hh signaling and that neither Bnl nor Hh signaling is sufficient (Fig. 7E). The conversion of Ci to its activator form by Hh signal transduction likely indicates that direct combinatorial regulation of *ct* and *aop* by Ci^Act^ and Pointed, which mediates transcription output of Bnl signaling, is unlikely. Further analysis of the elements that regulate *ct* and *aop* will be necessary to understand the basis for the combinatorial regulation.

## Acknowledgements

We thank: K. Irvine, M. Gibson, and X. Zheng for antibodies and the Bloomington Stock Center and Vienna Drosophila Resource Center for fly stocks. This work was funded by NIH T32HL007185 to R.H and R35GM122548 to T.B.K.

## Materials and Method

### Fly lines

*disp*^-^ (Burke et al., 1999); *Gal4*/*LexA* lines: *btl-Gal4* (Sato and Kornberg, 2002), *hh-Gal4* (Tanimoto et al., 2000), *Ay-Gal4* (Bloomington#3953), *btl-LHG* (Roy et al., 2014), *1151-Gal4* (Roy and VijayRaghavan, 1997); *UAS* lines: *UAS-mcd8:GFP* (Roy et al., 2011), *UAS-mcd8:Cherry* (Roy et al., 2011), *UAS-Ptc* (Johnson et al., 1995), *UAS-Ptc*^□loop2^ (Briscoe et al., 2001), *UAS-SmoRNAi* (VDRC#9542), *UAS–enRNAi* (Bloomington #33715), *UAS-Hh:GFP* (Torroja et al., 2004), *UAS-CD4:mIFP* (Yu et al., 2015), *UAS-CD4:GFP1-10* (from Kristen Scott), *UAS-SCAR-RNAi* (VDRC#21908), *UAS-disp RNAi* (Bloomington#27247), *UAS-dia-RNAi* (Bloomington), *UAS-hhRNAi* (Bloomington#31475), *UAS-Btl:Cherry* (Roy et al., 2011), *UAS-Ptc:GFP* (Torroja et al., 2004); *LexO* lines: *lexO-CD2:GFP* (Torroja et al., 2004), *lexO-CD4:GFP*^*11*^ (Roy et al., 2014), *lexO-n-syb-GFP*^*11*^ (Macpherson et al., 2015), *lexO-diaRNAi* (Fereres et al., 2019), *lexO-cherry:CAAX* (from K. Basler); *BAC*/*Crispr*/*Fosmid* Knock-in lines: *Ptc:Cherry BAC* (Chen et al., 2017); *Smo:GFP* BAC (Chen et al., 2017); *Ci:Cherry* (this study), *Cact:GFP* fosmid (VDRC#318145); *Hh:GFP* BAC (Chen et al., 2017), Btl:Cherry (Du et al., 2018); Enhancer traps: *hh-lacZ* (Lee et al., 1992), *ptc:GFP* (Bloomington#65320).

All crosses were at 25°C except those involving temporal induction of gene expression using *Gal80*^*ts*^. For cytoneme ablation experiments, *Gal80*^*ts*^ was done according to Roy et al., 2014. For overexpression of Hh:CD2, larvae were grown at 18°C to L3 incubated at 29°C for 24 hours.

### Generation of Cherry:Ci line

Using CRISPR-Cas9 gene editing, mCherry together with a short linker was inserted at the N terminus of the ci coding sequence. The template vector contains a left homology arm, mCherry, and right homology arm that were amplified by PCR. Left homology arm contains an overlapping sequence with approximately 1kb sequence upstream of the transcriptional start site. The right homology arm fragment contains an overlapping sequence with approximately 1kb sequence from the transcriptional start site. The three fragments were joined and cloned into PBS-SK using Gibson Assembly (NEB). The resulting vector is designated as Cherry:Ci donor vector. gRNA was cloned into pCDF3 (Addgene).

The following primers were used:

L-arm-fwd: cggtatcgataagcttgatGTTTTGCGCTGTTTGTGGACGTTAGAGTG

L-arm-rev: GGACGAGCTGTACAAGTCATGGACTAACTTTAATGAAATGGACGCGTACGCG

Cherry fwd: CATTAAAGTTAGTCCATGACTTGTACAGCTCGTCCATGCCGC Cherry rev: CGACGTCATTCTTGTTGATGGTGAGCAAGGGCGAGGAGGATAAC

R-arm-fwd: CTCGCCCTTGCTCACCATCAACAAGAATGACGTCGTTTTATAAATTC

R-arm-rev: ccgggctgcaggaattcgatGGCGTTGCCAATAACTTTTGCG

gRNA sequence: AAAATATGTAGGTAACGCGTAGG

### Immunohistochemistry and fluorescent imaging

L3 larvae were dissected in PBS; wing discs together with the Tr2 trachea were fixed in 4% formaldehyde in PBS. Tissue was washed in PBS-TritonX-100 (0.3%) followed by Roche Blocking Solution. The following antibodies were used: α-Dpp-prodomain (M. Gibson), mouse α-GFP (Roche), rabbit α-RFP (Rockland), mouse α-Ptc (DSHB, Apa1), mouse α-En (DSHB, 4D9), DAPI, Alexa633 conjugated-Phalloidin (Invitrogen), mouse α-Cut (DSHB, 2B10), mouse α-Aop (8B12H9), rat α-Ser (from Kenneth Irvine), rat α-Ihog (from Xiaoyan Zheng), rat α-Ci-AbN. Secondary antibodies from Invitrogen were used and samples were mounted in Vectashield (Vector labs).

Unfixed wing discs and ASPs were observed using the hanging drop method (Huang and Kornberg, 2016). All images were taken using the FV3000 Olympus Confocal microscope with GaAsP PMT detectors. Images were analyzed and processed with ImageJ and Photoshop.

### Analysis of the number of Hh:GFP puncta in the ASP

Maximum projections of 15 optical sections that span the upper and lower layers of the ASP were processed in ImageJ. For Fig. 1H, the position of each puncta was measured relative to the distance from the tip of the ASP. Histograms show the number of puncta in different bins at distances from the tip of the ASP. Fig. 5Q: the total number of Hh:GFP puncta in the ASP was tabulated.

### Analysis of cytoneme number and length

*btl-Gal4, UAS-mCD8:GFP/+; Gal80ts/+* (Control), *btl-Gal4, UAS-mCD8:GFP/UAS-Ptc*; *Gal80ts/+* (Ptc overexpression), *btl-Gal4, UAS-mCD8:GFP/+; Gal80ts/UAS-Hh:CD2* (Hh:CD2 overexpression) larvae were grown at 18°C until early L3 larvae and were incubated at 29°C for 24 hours prior to dissection. ASP cytonemes were imaged using the hanging drop method (Huang and Kornberg, 2016). For the analysis of cytoneme density, the number of cytonemes in the ASP bulb was counted and the number is expressed relative to perimeter length. The length and location of base was measured in ImageJ for each cytoneme. Cytonemes were categorized into tip (≤25μm from tip) and medial (>25μm from tip) based on the position of their base with respect to the tip of the ASP. The fraction of cytonemes contained in various length bins was measured for each sample. 4-6 samples were analyzed for each genotype.

### Area measurement

For the area of the entire wing disc and the entire notum, maximum projection of wing discs were created and the areas were measured in ImageJ. A 150µm x 150µm boxed region in the notum dorsal to the hinge/notum fold containing the band of Ptc expressing cells in the notum is defined as the microenvironment. Areas of myoblast, ASP, and notum in the microenvironment, or area of α-Ptc staining area within the myoblast, ASP, or notum were calculated in ImageJ from single optical sections.

### En intensity measurements in the ASP

Average projection of 15 optical sections was calculated in ImageJ for optical sections spanning the upper and lower layer of the ASP. α-En staining intensity was measured inside the ASP in these images and normalized to α-En staining intensity in the notum.

### Analysis of the morphology of myoblast layers

In ImageJ, orthogonal sections were generated along the region of the notum shown in Fig. 3A; depths of the myoblast layers were measured from the basal side of the wing discs.

