## Supplemental Figure 1 for "Hedgehog produced by the Drosophila wing imaginal disc induces distinct expression responses in three target tissues"

### Figure S1

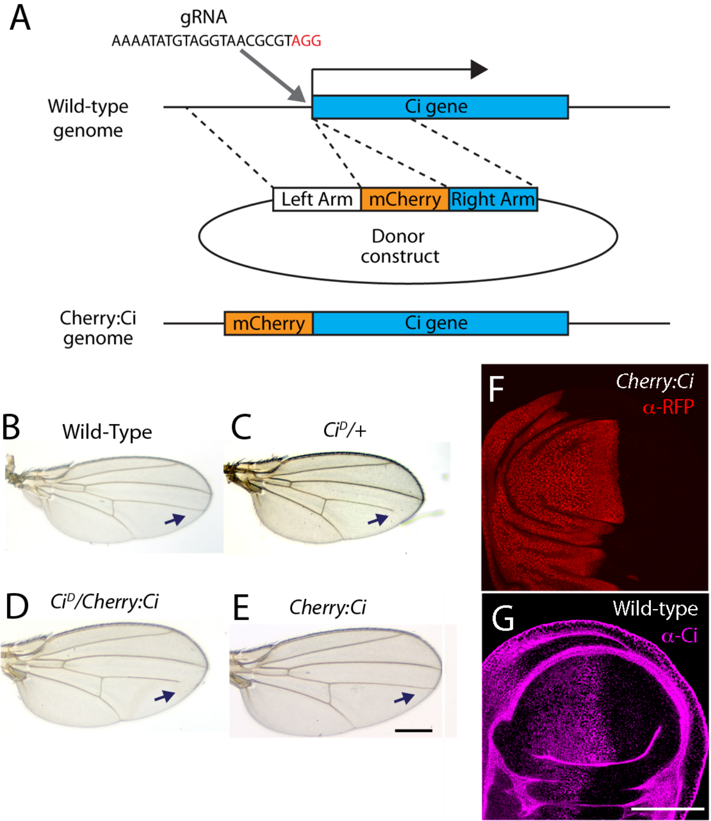
A
gRNA
AAAATATGTAGGTAACGCGTAGG
Wild-type genome
Ci gene
Left Arm
mCherry
Right Arm
Donor construct
Cherry:Ci genome
mCherry
Ci gene
B
Wild-Type
C
Ci^D/+
D
Ci^D/Cherry:Ci
E
Cherry:Ci
F
Cherry:Ci
α-RFP
G
Wild-type
α-Ci

**Figure S1. Design of Cherry:Ci Crispr** (A) Schematic showing the design of the Cherry:Ci CRISPR knock-in. (B-E) Adult wings of indicated genotype. Arrows indicate the termini of wing vein L4. Scale is 1μm. (F,G) Frontal views of wild-type and Cherry:Ci expressing wing discs stained with α-RFP or α-CiAbN, respectively. Scale bar is 50μm.
